# Population genomics of *Salmonella* enterica serovar Weltevreden ST365, an emerging predominant causative agent of diarrheal disease

**DOI:** 10.1101/2021.04.15.440096

**Authors:** Jianmin Zhang, Zhong Peng, Kaifeng Chen, Zeqiang Zhan, Haiyan Shen, Saixiang Feng, Hongchao Gou, Xiaoyun Qu, Mark Ziemann, Daniel S. Layton, Bin Wu, Xuebin Xu, Ming Liao

**Affiliations:** National and Regional Joint Engineering Laboratory for Medicament of Zoonoses Prevention and Control; Key Laboratory of Zoonoses, Ministry of Agriculture; Key Laboratory of Zoonoses Prevention and Control of Guangdong Province; Animal Infectious Diseases Laboratory, College of Veterinary Medicine, South China Agricultural University, Guangzhou 510642, China; State Key Laboratory of Agricultural Microbiology; The Cooperative Innovation Center for Sustainable Pig Production; College of Veterinary Medicne, Huazhong Agricultural University, Wuhan 430070, China; Institute of Animal Health, Guangdong Academy of Agricultural Sciences, Guangzhou 510640, China; School of Life and Environmental Sciences, Deakin University, Waurn Ponds Campus, Geelong, VIC 3215, Australia; Commonwealth Scientific and Industrial Research Organization Health and Biosecurity, Australian Centre for Disease Prevention, East Geelong, VIC 3220, Australia; Shanghai Municipal Center for Disease Control and Prevention, Shanghai 200336, China

**Author notes:** Address correspondence to Ming Liao,; Xuebin Xu,; or Bin Wu. Jianmin Zhang and Zhong Peng contributed equally to this work. Author order was determined on the basis of seniority.

**Keywords:** *Salmonella* Weltevreden, ST365, Population genomics, Antimicrobial resistance, Virulence factors encoding genes, Virulence plasmid

## Abstract

*Salmonella* enterica serovar Weltevreden is a recently emerged pathogen, and as such we lack a comprehensive knowledge of its microbiology, genomics, epidemiology and biogeography. In this study, we analyzed 174 novel *S*. Weltevreden isolates including 111 isolates recovered from diarrheal patients in China between 2006 and 2017. Our results demonstrate that the ST365 clone was the predominant causative agent of the diarrhea-outbreak during this period, as vast majority of the isolates recovered from diarrheal patients belonged to this sequence type (97.37%, 74/76). We also determined the ST365 clone as the predominant sequence type of *S*. Weltevreden from diarrheal patients globally from previously published sequences (97.51%, 196/201). In order to determine the possible antimicrobial genes and virulence factors associated with *S*. Weltevreden, we performed whole genome sequencing on our novel isolates. We were able to identify a range of key virulence factors associated with *S*. Weltevreden that are likely to be beneficial to their fitness and pathogenesis. Furthermore, we were able to isolate a novel 100.03-kb IncFII(S) type virulence plasmid that used the same replicon as pSPCV virulence plasmid. Importantly, we demonstrated through plasmid elimination a functional role for this plasmid in bacterial virulence. These findings are critical to further our knowledge of this high consequence pathogen.

**Importance:** *Salmonella* Weltevreden is a newly emerged foodborne pathogen and has caused several outbreaks of diarrheal diseases in some regions in the world. However, comprehensive knowledge of microbiology, genomics, epidemiology and biogeography of this newly emerged pathogen is still lack. In this study, we made an unexpected discovery that *S*. Weltevreden sequence type (ST) 365 is the causative agent in the diarrhea-outbreak in China and many other regions of the world. We also shown that this sequence type was widely recovered from animal, food, and environmental samples collected in different regions in the world. Importantly, we discovered a novel IncFII(S) type virulence plasmid commonly carried by *S*. Weltevreden strains of both human, animal, and food origins. These data facilitate future studies investigating the emergence of *S*. Weltevreden involved in diarrheal outbreaks and the global spread of *S*. Weltevreden strains.

## Introduction

*Salmonella* is a key global cause of human diarrheal diseases, and Salmonellosis is the third leading cause of death among the diarrheal diseases worldwide(1, 2). According to the Centers for Disease Control and Prevention, *Salmonella* bacteria cause approximately 1.35 million infections, 26,500 hospitalizations, and 420 deaths in the United States every year(3). Consumption of contaminated food, particularly food of animal origin, such as eggs, meat, poultry, and milk are the main source of *Salmonella* infections. This is due to the high prevalence of *Salmonella* bacteria in animals, particularly in food animals such as poultry, pigs, and cattle. *Salmonella* can pass through the entire food chain from animal feed, primary production, as well as all the way to households or food-service establishments and institutions(1, 4).

To date, *Salmonella* bacteria are classified into six subspecies containing over 2500 serovars(5). Among these serovars, *S*. enterica serovar Weltevreden has been recognized as a newly emerged pathogen and has caused several outbreaks in different regions in the world, including Réunion Island (6), Europe (e.g. Norway, Denmark and Finland) (7), and Southeast Asia (e.g. India, Malaysia, Thailand, Laos)(8–14). Recently, a foodborne outbreak of *S*. Weltevreden sequence type (ST) 1500 caused an acute watery diarrheal illness in 150 students aged between 20~30 years in Pune, India(15). These reports suggest *S*. Weltevreden represents a significant threat to global public health, however, there is still a lack of genomic characterization and assessment of virulence of *S*. Weltevreden remain limited (16). In this study, we report the epidemiological distribution, the microbiological and genomic characteristics, as well as the virulence of *S*. Weltevreden strains globally.

## Results

### *S.* Weltevreden ST365 originated from contaminated food is likely to be responsible for the outbreak of human diarrhea in four provinces in China between 2006 and 2017

Between 2006 and 2017, we recorded 111 cases of diarrhea in Guangdong, Guangxi, Shanghai, and Yunnan provinces in China (Fig. 1A; Supplementary materials Table S1). In each of the patients, *S*. Weltevreden strains were recovered from the stool or blood samples (Fig. 1B; Supplementary materials Table S1). To understand the genomic characteristics of these *S*. Weltevreden isolates, we randomly selected 76 strains (76/111) for Illumina sequencing (Supplementary materials Table S2). Multilocus sequence typing (MLST) analysis revealed the majority of the isolates typed belonged to ST365 (74/76) (Fig. 1C). The remaining two isolates belonged to ST155 (1/76) and ST648 (1/76), which were responsible for two human diarrheal diseases in Shanghai.

**Fig. 1.**
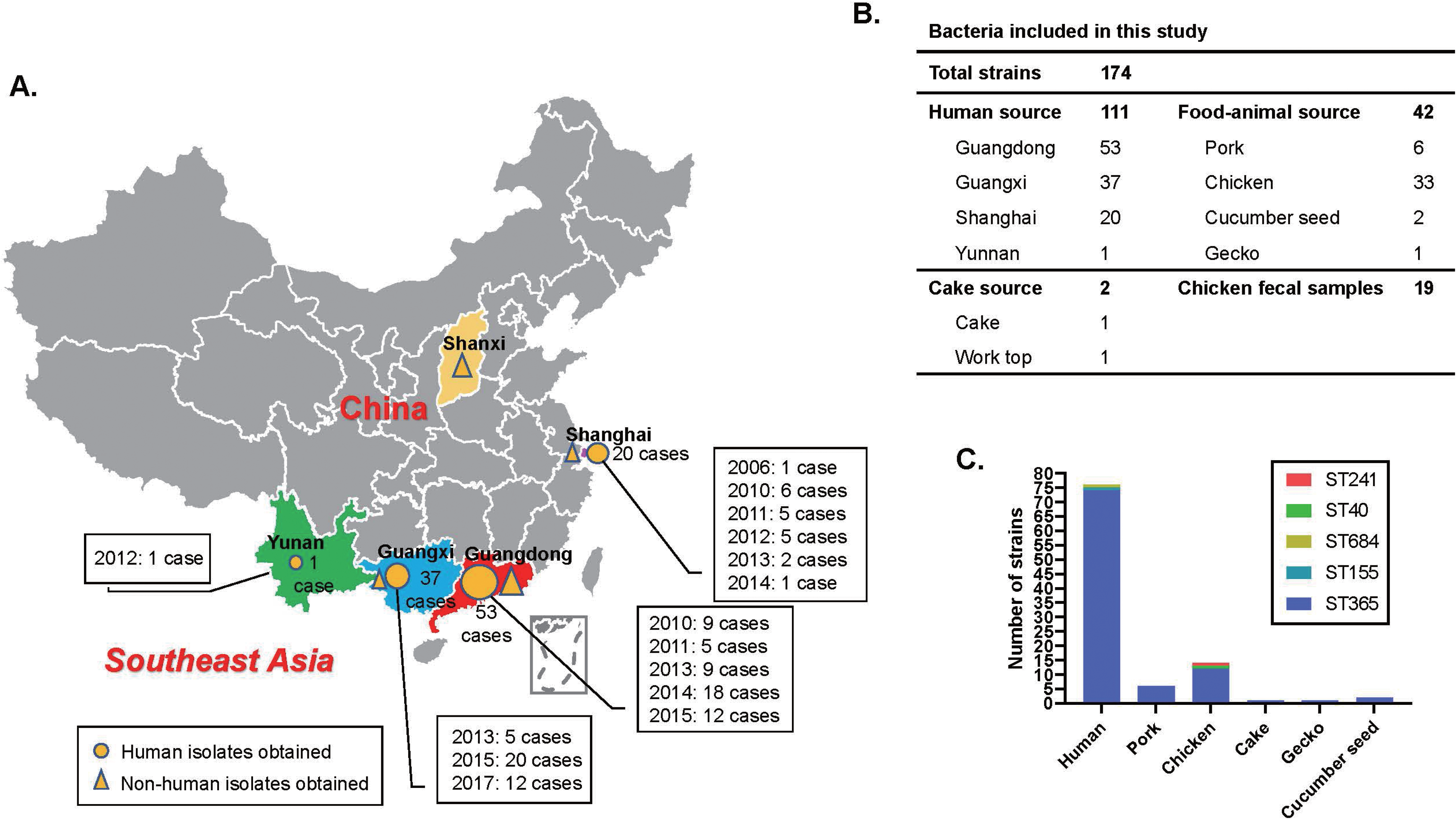
Geographical distribution and sequence type analysis of *S*. Weltevreden isolates. **(A.)** Shows the temporal and geographical location of human *S*. Weltevreden infections in China; **(B.)** Show the breakdown of *S*. Weltevreden isolates by origin used in this study; **(C.)** The bar graph demonstrates the sequence types of *S*. Weltevreden isolates from different sources.

Our questionnaire survey revealed that most of the patients had an exposure to chicken, however several were exposure to pork, seafood, or cake prior to presenting with symptoms of fever, emesis, diarrhea, and/or stomach ache. These information suggest contaminated food might be a source for the dissemination of *S*. Weltevreden to humans and lead to the diarrhea. To verify this hypothesis, we studied 63 *S*. Weltevreden strains from different types of food associated samples (pork, poultry, seafood, cake), as well as other animals and environmental samples collected between the same time period (Fig. 1B; Supplementary materials Table S1). Our PFGE typing revealed most of the human *S*. Weltevreden isolates were the same PFGE types with those isolates from chicken and/or chicken feces collected from Guangdong, and several human isolates shared the same types with those strains recovered from pork or cucumber seed (Fig. 2). We also randomly selected 24 strains from these 63 isolates for Illumina sequencing (Supplementary materials Table S2). Strikingly, MLST analysis revealed that the majority of the isolates belonged ST365 (22/24) (Fig. 1C). The remaining two belonged to ST40 (1/24) and ST241 (1/24) (Fig. 1C).

**Fig. 2.**
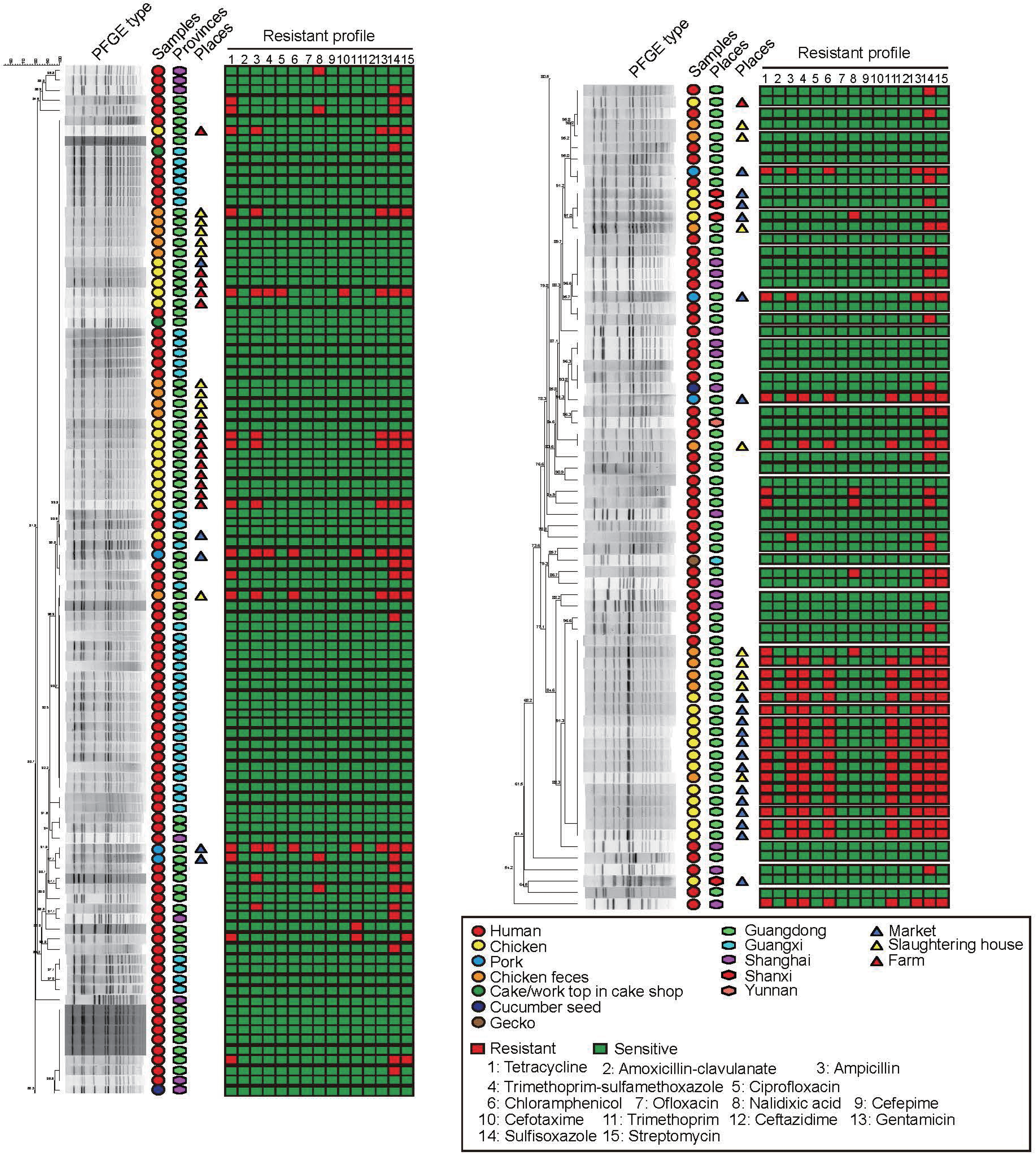
PFGE patterns of *S*. Weltevreden strains with different sources of China. The heat map shows the resistance profiles of individual isolates. Red indicates resistance and green indicates sensitivity. Geographic regions of the isolates are marked with hexagons; hosts are marked with circles; triangles indicate the places of non-human originated isolates.

### *S*. Weltevreden ST365 is widely recovered from diarrheal humans and food-associated samples in the world

To further investigate the prevalence of *S*. Weltevreden ST365, we next studied the whole genome sequences of 178 genome sequences of *S*. Weltevreden publicly available in NCBI as of 31 August 2020 (Supplementary materials Table S3). According to the biosample information registered, the sequences were derived from diarrheal humans (125/178), animals (15/178), food including meat, seafood, vegetable, and the other types of food (24/178), environment samples (4/178), and porcine feed (2/178) in Asia, Europe, North America, Africa, South America, and Oceanian. The remaining eight strains lack information of host type and places of isolation. Determination of the sequence types revealed that again the vast majority of these *S*. Weltevreden isolates belonged to ST365 (170/178); the other determined sequence types included ST3771 (3/178), ST2183 (1/178), ST2383 (1/178), ST3902 (1/178), and two novel sequence types (Fig. 3; Supplementary materials Table S3). Among the 125 human isolates, with the exception of only 3 isolates from Sri Lanka determined as ST3771, all of the remaining 122 isolates were ST365 (Supplementary materials Table S3). *S*. Weltevreden ST365 was prevalent in many regions in Asia, Africa, Europe, North America, and Oceanian (Fig. 3).

**Fig. 3.**
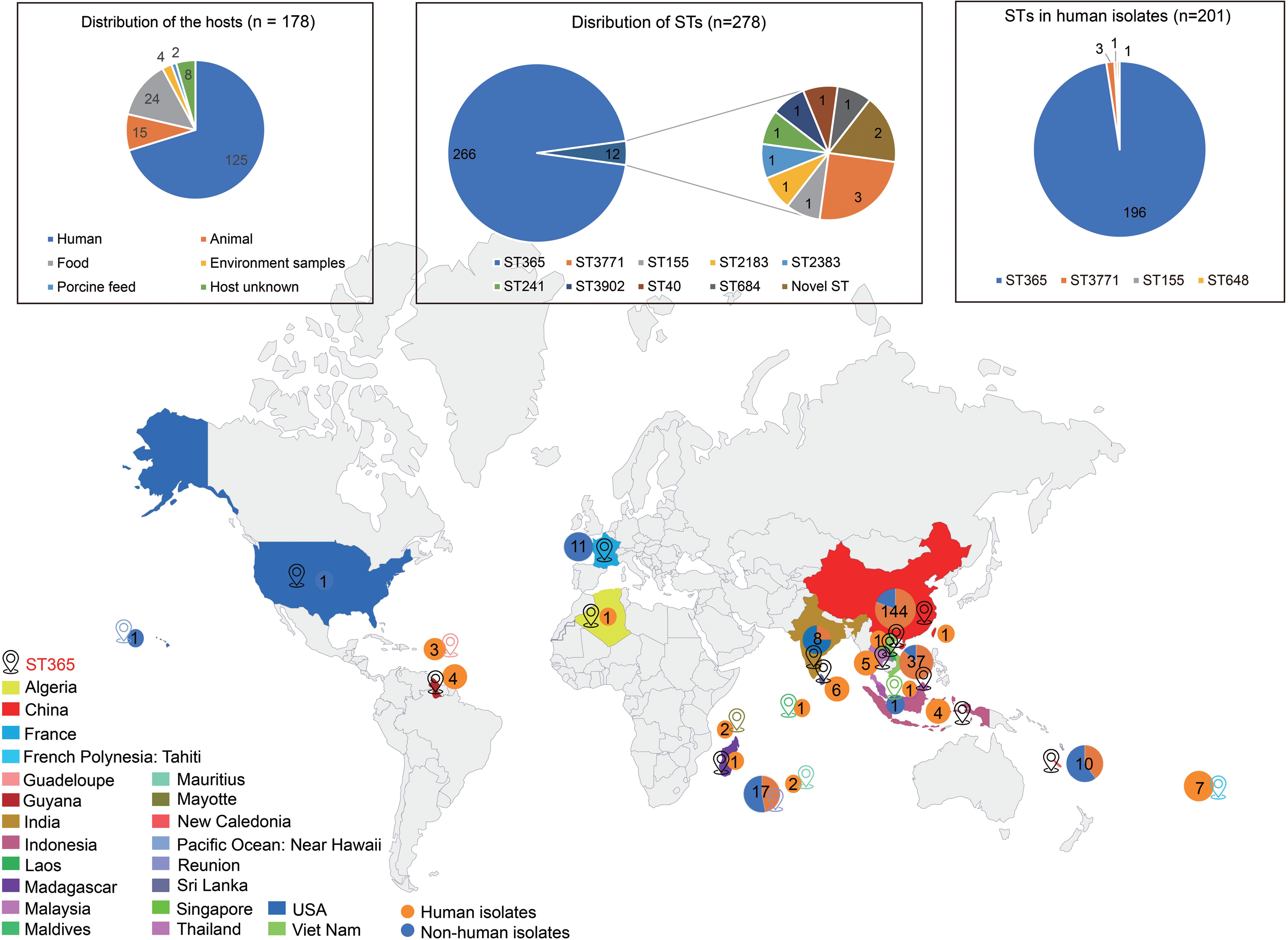
Global distribution of *S*. Weltevreden ST365. Orange circles indicate the countries/regions where *S*. Weltevreden isolates were obtained. Numbers in the circles refer to the numbers of isolates from each of the countries or regions. Summaries of the host information of the 178 isolates from NCBI as well as sequence types of all 278 isolates including 100 novel isolates sequenced in this study, and sequence types of human isolates are shown at the top left corner, top center, and top right corner, respectively.

Phylogenetic analysis revealed that *S*. Weltevreden strains associated with human diarrhea and/or those isolated from food animals, seafood, as well as environmental samples in China displayed a close relationship with those recovered from diarrheal patients and different food types outside China (Fig. 4). Interestingly, those non-ST365 strains (including ST40, ST155, ST241, ST684, ST2183, ST2383, ST3771, and ST3902) demonstrated a close phylogenetic relationship with the ST365 strains (Fig. 4). According to the Enterobase *Salmonella* MLST Database, ST365, ST2183, ST2383, ST3771, and ST3902 were assigned into the Clonal complex 205, while ST40, ST155, ST241, ST684 were assigned into Clonal complex 57, 237, 33, and 157, respectively. These results indicate a strong relationship between the ST365 clone and the presence of diarrheal disease in humans.

**Fig. 4.**
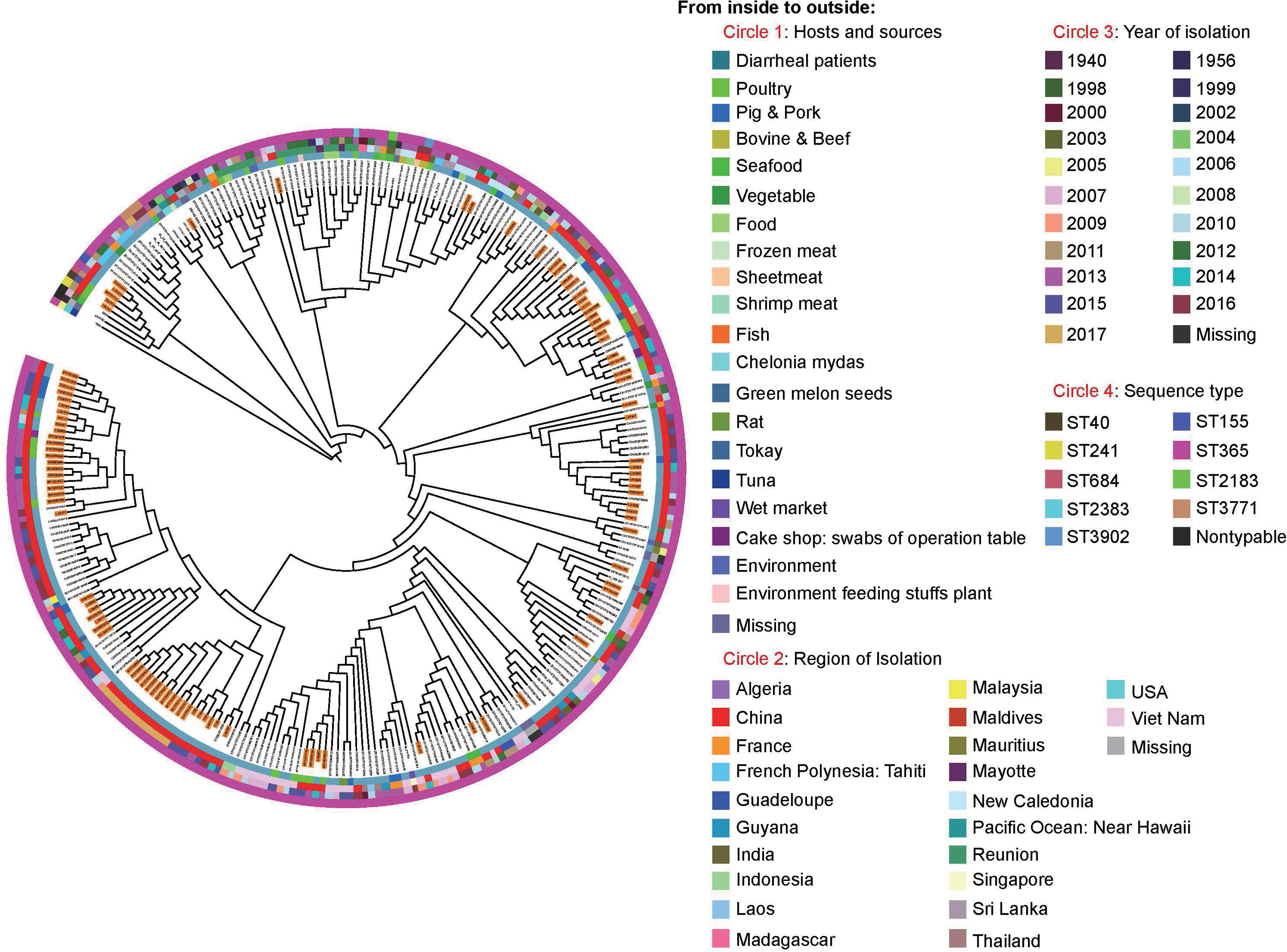
Phylogenetic relationship of *S*. Weltevreden isolates from different regions, hosts and sequence types. The tree was generated based on the single nucleotide polymorphisms across the whole genome sequence (gSNPs) by using the PHYLIP (version 3.698) software. Circles from inside to outside indicate the hosts of the isolates (circle 1), regions of isolation (circle 2), years of isolation (circle 3), and the sequence types of the isolates (circle 4), respectively. Isolates with background highlighted in orange on the tree are those sequenced in the present study.

### *S*. Weltevreden ST365 does not shown severe antimicrobial resistance profile

To further understand the *S*. Weltevreden serovar, in particular the ST365 clone, we performed prediction of ARGs using the 178 whole genome sequences. Interestingly, the *S*. Weltevreden strains did not contain a particular abundance of ARGs (Fig. 5; Supplementary materials Table S4) indicating that the antimicrobial resistance (AMR) profiles of *S*. Weltevreden may not be of serious concern. However, those ARGs contained may confer the bacteria resistance to antimicrobials belonging to aminoglycosides, rifampicin, beta-lactams, phenicols, trimethoprim, Macrolide-Lincosamide-Streptogramin B, fosfomycin, colistin, fluoroquinolones, sulphonamides, and tetracyclines (Fig. 5). To further investigate this, we tested the susceptibility of the 111 novel *S*. Weltevreden isolates including 96 *S*. Weltevreden ST365 we collected between 2006 and 2017 on 15 types of antibiotics belonging to the above classes. In agreement with the prediction of ARGs, the antimicrobial susceptibility testing (AST) results revealed that 111 *S*. Weltevreden strains including the 96 *S*. Weltevreden ST365 were susceptible to many types of antimicrobials tested, and the carried ARGs conferred the isolates resistance to the corresponding antimicrobials (Fig. 2).

**Fig. 5.**
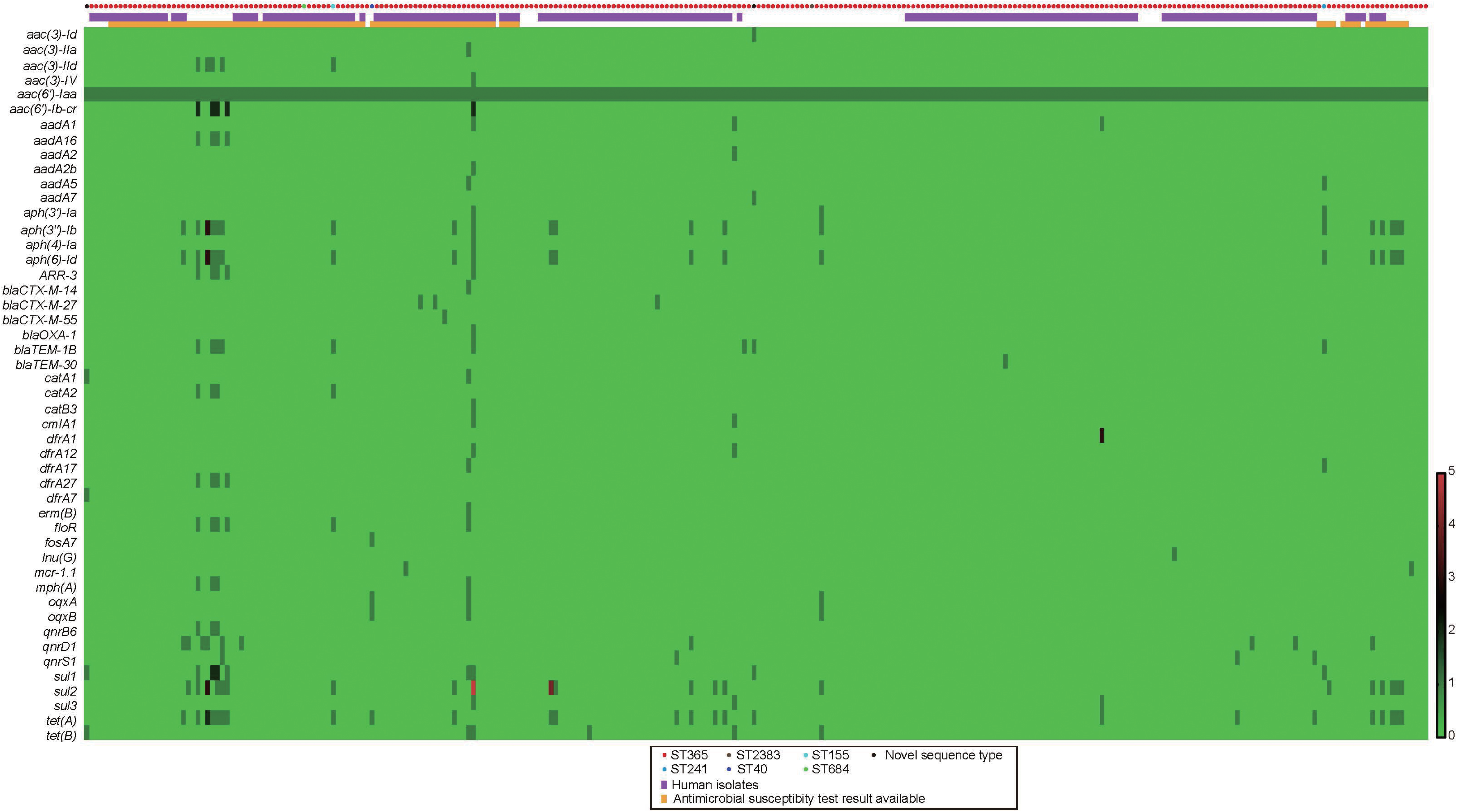
Antimicrobial resistance genes carried by 278 *S*. Weltevreden isolates. The number of antimicrobial resistance gene in each of the isolates are shown with bars at lower right corner. Colored circles refer to the sequence types of different isolates. Orange rectangles show isolates with antimicrobial susceptibility test results available (See Figure 2). Purple rectangles show isolates from humans. Isolates’ names (from left to right) are provided in Supplementary materials Table S4.

### A novel T4SS-carrying-IncFII(S) type plasmid is found to be associated with virulency

To understand the genomic basis for pathogenesis, we first determined the VFGs carried by *S*. Weltevreden. This strategy identified a total of 558 types of VFGs (Supplementary materials Table S5). These VFGs encoded proteins participating in bacterial adherence (e.g., Lpf, MisL, RatB, ShdA, SinH, type 1 fimbriae), magnesium uptake (e.g., MgtBC), resistance to antimicrobial peptides (e.g., Mig-14), serum resistance (e.g., Rck), anti-stress (e.g., SodCI) and toxin (e.g, typhoid toxin CdtB). Among these 558 types of VFGs, 441 were were carried by more than 90% of the *S*. Weltevreden (*n* = 251). Of particular note were *fbpC*, *hitC*, *cylA*, *ptxR*, *phoP*, *cdpA*, *phoR*, *bfmR*, *lap*, *bauE*, *pilW*, *chuV*, and *mgtB*; as more than five copies of these VFGs were found in each of the *S*. Weltevreden isolates (Supplementary materials Table S5).

We next aimed to determined putative plasmids carried by these 278 S. Weltevreden isolates and identified a total of 21 different types of plasmid replicons (Fig. 6A; Supplementary materials Table S6). Interestingly, among these plasmid replicons, an IncFII(S) type plasmid was present in 78.75% of the *S*. Weltevreden strains (226/278; Fig. 6A). This plasmid had the same replicon as the virulence plasmid pSPCV (GenBank accession number: CP000858) which shares very high sequence identity with the *S*. typhimurium virulence plasmids pSLT(17). We did not however, observe pSPCV or pSLT homologous sequences in the genome sequences of the 278 *S*. Weltevreden isolates. To further investigate this novel plasmid, we generated the complete genome sequences of the IncFII(S) type plasmid (designated pSH17G0407; GenBank accession no. MW405382) harbored in isolate SH17G0407 using ONT sequencing. The strategy yielded a plasmid of 100.03-kb in size with a G+C content of approximately 49.2% (Fig. 6B). This plasmid showed phylogenetic relatedness to the *Salmonella* virulence plasmids pSPCV (Fig. 6C). Of high importance, we identified a putative T4SS encoding region in pSH17G0407. This region contained 63 genes and was flanked by two insertion sequences belonging to the IS*256* family (ISSod4) and the IS*4* family (ISSfl1) (Fig. 6B). At least 28 genes in this region encoded proteins involved in the composition of a putative T4SS (Fig. 6B). In addition, this region also contained many genes encoding proteins involved in DNA replication, mobility, and conjugation (Fig. 6B). However, it remains unclear whether this region represents a single transposable unit or a mosaic of gene acquisition events in the plasmid. Notably, a homologous sequence of pSH17G0407 was present in the genomes of the 266 *S*. Weltevreden strains (Supplementary materials Table S7) suggesting it is of high functional importance to ST365 clone.

**Fig. 6.**
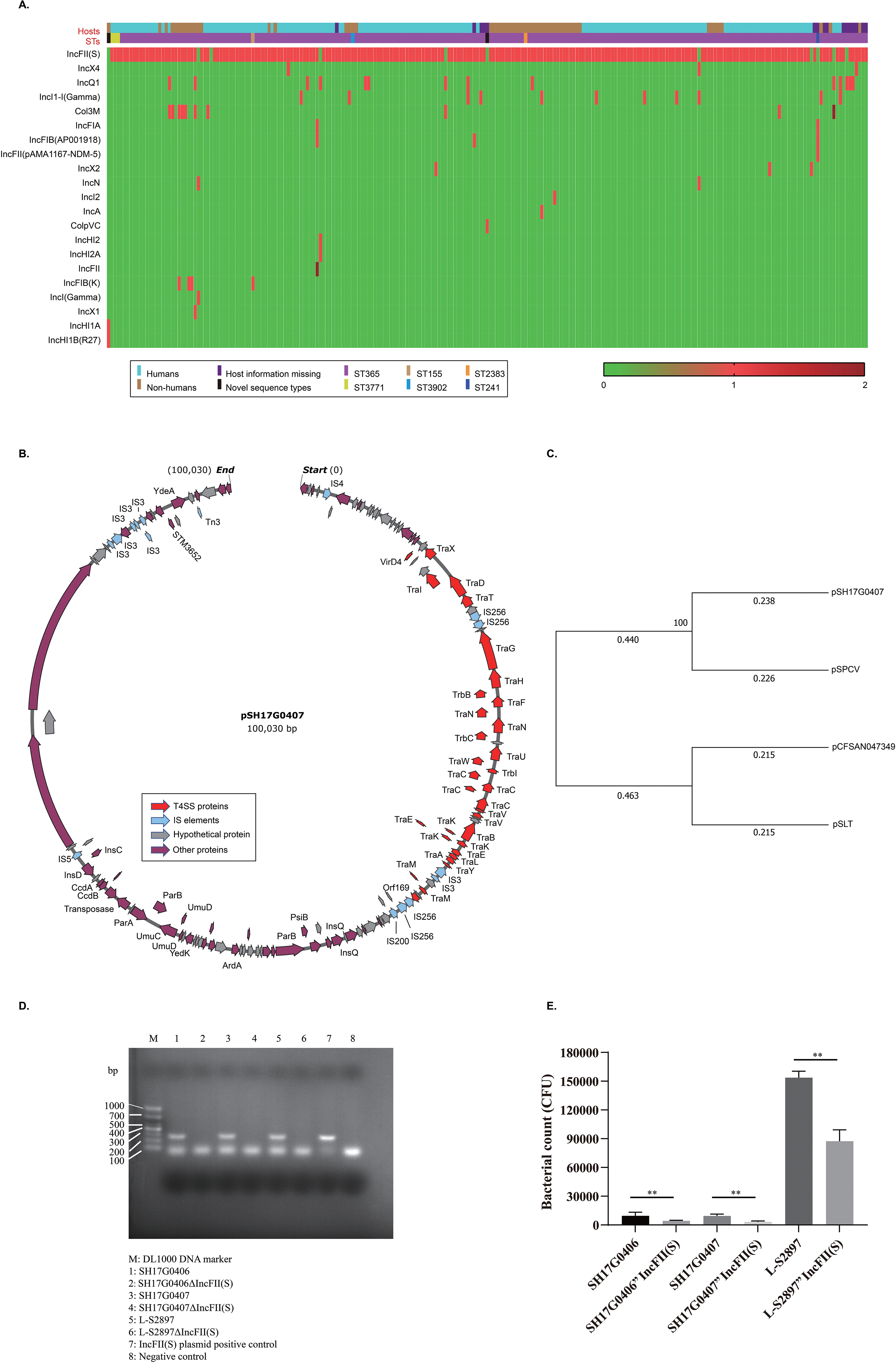
Heatmap showing putative plasmids carried by 237 *S*. Weltevreden isolates. **(A.)** The number of plasmid replicons in each of the isolates are shown with bars at lower right corner. Rectangles in different colors show the sequence types and hosts information, which are given at lower left corner. Isolates’ names (from left to right) are provided in Table S6 in supplementary materials. **(B.)** Map of plasmid pSH17G0407 from isolate SH17G0407. Predicted coding sequences were shown used arrows in different colors (grey: genes encoding hypothetic proteins; red: genes encoding T4SS proteins). **(C.)** Phylogenetic relationships of plasmid pSH17G0407 and *Salmonella* virulence plasmids pSPCV (GenBank accession number: CP000858), pSLT (GenBank accession number: LN999012), and pCFSAN047349 (GenBank accession number: CP040702). **(D.)** PCR results indicating the elimination of the T4SS-carrying-IncFII(S) type plasmid. **(E.)** The number of *S*. Weltevreden strains and their IncFII(S)-plasmid elimination strains invading to HeLa cells. Data represents mean ± SD. The significance level was set at *P* < 0.05 (*) or *P* < 0.01 (**).

To determine the extent of virulence conferred by this plasmid we performed plasmid elimination experiments to study the influence of the T4SS-carrying-IncFII(S) type plasmid on virulence of *S*. Weltevreden. We eliminated these plasmids in three *S*. Weltevreden isolates (SH17G0406, SH17G0407, and L-S2897) (Fig. 6D). Comparisons of bacterial invasion to HeLa cells between the wild type strains (SH17G0406, SH17G0407, and L-S2897) and plasmid-curing strains (SH17G0406ΔIncFII(S), SH17G0407ΔIncFII(S), and L-S2897ΔIncFII(S)) revealed that the elimination of the plasmid significantly decreased the bacterial invasion to the cells (Fig. 6E), suggesting this plasmid is important for the bacterial virulence.

## Discussion

In this study, we reported the distribution, microbiological and genomic characteristics of a newly emerged diarrhea associated *Salmonella* serovar named Weltevreden isolated from different regions around the world. In China, a recent study has reported an outbreak of *S*. Weltevreden infections in Guangdong province between 2015 and 2016(8). While *S*. Weltevreden has been detected in the poultry supply chain and other meat samples in China previously(18, 19), outbreaks of human diarrhea caused by *S*. Weltevreden have not been reported until recently(8). Here, our retrospective study revealed that *S*. Weltevreden associated human diarrhea occurred in many other parts in China in addition to Guangdong between 2006 and 2017 (Fig. 1). It has been reported that consumption of contaminated foods and seafood is recognized as the main cause of *S*. Weltevreden infection in humans(2). Consistently, our PFGE typing results showed that *S*. Weltevreden strains recovered from either stool samples or blood of the patients during the outbreaks in China between 2006 and 2017 had similar PFGE patterns with those isolated from poultry, pork, and/or other food types in south China (Fig. 2). In particular, *S*. Weltevreden strains with similar PFGE types were also isolated from chicken/pig farms, slaughtering houses, and markets (Fig. 2). These findings suggest that animals, particularly food animals and their products are an important source of the spread of *S*. Weltevreden to humans.

In a previous report of human diarrhea caused by *S*. Weltevreden in Guangdong in China (8), the ST365 clone was found to be responsible for this outbreak. Apart from this report, there is no reports of the outbreak of this clone in the other regions in China or globally^1^. Here, we demonstrated that the ST365 clone was widely recovered from human diarrheal cases as well as from both animal and environmental samples in different parts of the world by analyzing the whole genome sequences (Figs 1C, 3, and 4). These findings indicate that *S*. Weltevreden ST365 is a worldwide pathogenic clone and represents high risks to human health.

Knowledge of the genetic mechanisms of AMR is critical for defining appropriate treatments, refining diagnostics, and conducting epidemiological studies of AMR (20). Interestingly, our prediction of ARGs and AMR phenotype determination revealed that *S*. Weltevreden strains including the ST365 clone did not show severe resistance profile (Figs 2 and 5), suggesting that many antimicrobial agents may still effective for the treatment of infections caused by *S*. Weltevreden. However, multidrug-resistance phenotypes were also determined, particularly among those isolates recovered from slaughterhouses and markets (Fig. 2). These isolates possess a strong possibility for transmission to humans. By determination of the carried VFGs, we also demonstrated the genetic mechanisms of pathogenesis. Our results revealed that each of the *S*. Weltevreden isolates including the ST365 clone possessed numerous VFGs (Supplementary materials Table S5). These VFGs encoded proteins participating in bacterial adherence, magnesium uptake, resistance to antimicrobial peptides, serum resistance, anti-stress, toxin, etc. All of these bioactivities are beneficial for bacterial survival and fitness in hosts and therefore contribute to the pathogenesis (21).

Mobile genetic elements particularly plasmids play important roles in the dissemination of ARGs or VFGs in many bacterial species particularly in members belonging to *Enterobacteriaceae*(22, 23). Here, we analyzed putative plasmids harbored in *S*. Weltevreden including the ST365 outbreak clone and found the wide presence of an IncFII(S) type plasmid (Fig. 6A). We identified and determined the sequence of a IncFII(S) type plasmid associated with the ST365 outbreak clone and other *S*. Weltevreden strains (Fig. 6B). Although we did not find a high abundance of VFGs on the plasmid, we discovered a putative T4SS (Fig. 6B). An important role of T4SS in bacteria is to deliver DNA or proteins including virulence proteins and toxins to target cells(24). Considering *S*. Weltevreden strains harbored many virulence factors, the presence of a T4SS may help deliver these virulence factors to host cells and therefore contributes to the pathogenesis of *S*. Weltevreden. However, detailed functions of this T4SS as well as its plasmid payload need to be characterized. Our plasmid elimination and cell invasion assays revealed that elimination of the plasmid significantly decreased the bacterial invasion to the cells (Fig. 6E). Since bacterial invasion to host cells is an important step for bacterial infection and pathogenesis(25, 26), it can be concluded that the wide presence of this T4SS-carrying-IncFII(S) type plasmid in *S*. Weltevreden strains contributes to the bacterial virulence.

In conclusions, we reported the epidemiological distribution, the microbiological and genomic characteristics, as well as the virulence of *S*. Weltevreden strains in this study. By whole genome sequencing and genome analyses, we found that *S*. Weltevreden strains particularly the ST365 clone was responsible for the outbreak of human diarrhea in China between 2006 and 2017. We also revealed that this outbreak clone was widely recovered from diarrheal patients in many regions in the world, suggesting that ST365 might be a worldwide pathogenic clone and represents a severe threat to human health. Since *S*. Weltevreden strains and the ST365 outbreak clone have been also recovered from food animals, food, and environmental samples, improving food safety is necessary. In addition, we also revealed how AMR phenotypes and virulence are associated with the genomes of *S*. Weltevreden strains, and provided novel data to these medically important features. Our framework presented herein will facilitate future studies investigating the emergence of *S*. Weltevreden involved in diarrheal outbreaks and the global spread of *S*. Weltevreden strains.

## Materials and methods

### *Salmonella enterica* serovar Weltevreden strains and genome sequences

A collection of 174 novel *S*. Weltevreden isolates were used in this study, including 111 isolates recovered from human diarrheal cases recorded in Guangdong, Guangxi, Shanghai, and Yunnan provinces in China between 2006 and 2017, and 63 *S*. Weltevreden strains recovered from poultry (*n* = 52), pork (*n* = 6), cucumber seed (*n* = 2), gecko (*n* = 1), cake (*n* = 1), and work top of a cake shop (*n* = 1) in Guangdong (*n* = 55), Guangxi (*n* = 2), Shanghai (*n* = 2), and Shanxi (*n* = 4) provinces during the same time period (Supplementary materials Table S1). As of 31 August 2020, there are 178 genome sequences of *S*. Weltevreden publicly available at NCBI (https://www.ncbi.nlm.nih.gov/genome/browse/#!/prokaryotes/152/Salmonella%20Weltevreden). These genome sequences were downloaded and included for analysis in this study (Supplementary materials Table S3).

### Antimicrobial susceptibility testing

Bacterial antimicrobial resistance phenotypes were tested using the broth microdilution methods (CLSI document M31-S1). A total of 15 types of antimicrobials belonging to aminoglycosides (gentamicin), beta-lactams (amoxicillin-clavulanate, ampicillin, cefepime, cefotaxime, ceftazidime), phenicols (chloramphenicol), trimethoprim, Macrolide-Lincosamide-Streptogramins (streptomycin), fluoroquinolones (ciprofloxacin, ofloxacin, nalidixic acid), sulphonamides (trimethoprim-sulfamethoxazole, sulfisoxazole), and tetracyclines (tetracycline) were tested. Results were interpreted using the CLSI breakpoints (CLSI M100, 28th Edition). Each of the antibiotics was tested with three duplicates. For quality control *E. coli* ATCC 25922 was used.

### Pulsed-field Gel Electrophoresis

Pulsed-field Gel Electrophoresis (PFGE) was performed by following the standardized protocol used by PulsedNet participating laboratories (27). Briefly, genomic DNA of each of the isolates were digested using the restriction enzyme *Xba*I and was then analyzed using PFGE, as described previously (28). *Salmonella* H9812 was used as a standard control strain. A molecular Imager Gel Doc XR System Universal Hood II (Bio-Rad Laboratories, CA, USA) was used to generate the PFGE gel pictures. Results were analyzed using the Bionumerics software (Version 5.1; Applied-Maths, Sint-Martens-Latem, Belgium).

### Whole genome sequencing by Illumina and Oxford Nanopore Technologies

We randomly selected 100 isolates including 76 human isolates and 24 animal isolates collected in this study for Illumina sequencing (Supplementary materials Table S2). Genomic DNA was extracted from broth cultures using a commercial Bacteria DNA Kit (TIANGEN, Beijing, China), and was then analyzed by electrophoresis on a 1% agarose gel as well as a Qubit 2.0 (Thermo Scientific, Waltham, USA). DNA libraries were generated using a NEBNext Ultra^TM^ II DNA Library Prep Kit (NEB, Ipswich, USA) and were then sequenced on an Illumina NovaSeq 6000 platform (Illumina, San Diego, USA) at Novogene Co. LTD (Tianjin, China), using the pair-end 350 bp sequencing protocol. Raw reads with low quality were filtered as previously described (29). High-quality reads were *de novo* assembled using SPAdes (version 3.9.0) (30) to generate contigs.

In addition, the complete sequence of a plasmid presence in *S*. Weltevreden isolate SH17G0407 was generated using Oxford Nanopore technology (ONT) in combination with the Illumina technology. Plasmid DNA was extracted using the phenol-chloroform protocol combined with Phase Lock Gel tubes (Qiagen GmbH) and was detected by the agarose gel electrophoresis as well as quantified by Qubit® 2.0 (Thermo Scientific, Waltham, USA). Libraries for ONT sequencing were prepared using an SQK-LSK109 kit of Oxford Nanopore Technologies Company; while libraries for Illumina sequencing were prepared by using a NEBNext® Ultra™ DNA Library Prep Kit for Illumina (NEB, USA) following manufacturer’s instructions. Prepared DNA libraries were sequenced using Nanopore PromethION platform and Illumina NovaSeq PE150 at Novogene Co. LTD (Tianjin, China), respectively. ONT and Illumina short reads were finally assembled and combined using the Unicycler v0.4.4 software with default parameters.

### Bioinformatic analysis

Sequence types (STs) and their mutilocus sequence typing (MLST) clonal complexes were analyzed by submitting the whole genome sequence against the Enterobase *Salmonella* MLST Database (http://enterobase.warwick.ac.uk/species/index/senterica). Genome sequences were annotated by using the RAST server (31). Antimicrobial resistance genes (ARGs), virulence factors encoding genes (VFGs), and plasmid types were predicted using ResFinder 4.0 (32), VFanalyzer (33), and PlasmidFinder 2.1 (34), respectively. Evolutionary trees based on genomic single nucleotide polymorphism (gSNP) were constructed with the Maximum Likelihood method and Tamura-Nei model in MEGAX software with 1000 bootstrap values (35), and were visualized using the iTOL online tool (36). Presence of type IV secretion system (T4SS) proteins and insertion elements were determined using SecReT4 2.0 (37) and IS finder (38), respectively.

### Plasmid elimination and cell invasion assay

An IncFII(S) type plasmid determined in most of the *S*. Weltevreden strains in this study was eliminated by using the ethidium bromide (EB) protocol as described previously (39). Briefly, a small inoculum (approximately 10^4^ CFU/ml) of *S*. Weltevreden were grown in Luria Bertani (LB) broth (Sigma-Aldrich, MO, USA) containing 30 μg/ml EB until slight turbidity observed. Afterwards, bacterial culture with appropriate dilution was plated on LB agar and incubated at 37 °C for overnight. Single colonies growing on the agar plates were selected and the elimination of the plasmid was examined by using PCR with primers targeting the IncFII(S) type replicons (F: 5’-CTGTCGTAAGCTGATGGC-3’; R: 5’-CTCTGCCACAAACTTCAGC-3’). If the PCR result is still positive for the IncFII(S) plasmid replicons, the above-mentioned bacterial subculture in the EB-containing medium should be performed until the plasmid was eliminated successfully.

To facilitate the analyses of invasion assays, HeLa cells (human cervical carcinoma, ATCC® CCL-2™) were cultured in Dulbecco’s modified eagle medium (DMEM, Thermo Fisher) supplemented with 10% (*v*/*v*) heat- inactivated fetal bovine serum (Gibco). Cells were seeded into 12-well plates (10^6^ cells per well) and cultured overnight. For bacterial preparation, overnight culture of plasmid-elimination strains and their wild-type strains were transformed into fresh LB broth at 1: 100 (*v*/*v*) and were incubated at 37 °C to OD_600_ = 1.0. After centrifugation at 4℃, 6000 rpm for 5 min, bacterial pellets were harvested and were washed using PBS for three times, followed by resuspension in DMEM. Each well of the cells were inoculated with either the plasmid-elimination strains or the wild type strains at a multiplicity of infection (MOI) value of 1:100. After incubation at 37 °C for 2 hours, the cells were washed using PBS for three times to remove the dissociative bacteria. Gentamicin (100 mg/ml) were given and the cells were incubated at 37 °C for 1 hours to kill bacteria adhesion on cell surface. Thereafter, cells were lysed using Triton X-100 buffer. A series of 10-fold dilution were performed to the lysed cells using PBS and appropriate dilutions were plated on LB agars. The agar plates were cultured at 37 °C overnight for bacterial count. Statistics analysis was performed using the “Two-way ANOVA” strategy in GraphPad Prism8.0. Data represents mean ± SD. The significance level was set at *P* < 0.05 (*) or *P* < 0.01 (**).

## Data availability

Whole genome sequences of the 100 *S*. enterica serovar Weltevreden strains obtained in the present study were deposited in GenBank with a BioProject ID PRJNA673740. Accession numbers for each of the genome sequences deposited are listed in Supplementary materials Table S2. The complete genome sequence of the plasmid harbored in strain SH17G0407 was also deposited in GenBank under an accession no. MW405382.

## Supplementary materials

**Table S1.** *S*. Weltevreden strains used in this study and their characteristics.

**Table S2.** *S*. Weltevreden strains sequenced in this study.

**Table S3.** *S*. Weltevreden genome sequences downloaded from NCBI as of 31 August 2020.

**Table S4.** Antimicrobial resistance genes (ARGs) presence in *S*. Weltevreden strains.

**Table S5.** Virulence factors encoding genes (VFGs) presence in *S*. Weltevreden strains.

**Table S6.** Plasmid replicons presence in *S*. Weltevreden strains.

**Table S7.** BLAST results of pSH17G0407 genome sequence against the 278 *S*. Weltevreden genome sequences.

## Acknowledgements

The authors thank staffs at Novogene Co. LTD (Tianjin, China) for technical support to perform Illumina and Oxford Nanopore sequencing. This work was supported by the National Key R&D Program of China (2017YFC1600101, 2018YFD0500500); National Natural Science Foundation of China (31972762); Guangdong Province Universities and Colleges Pearl River Scholar Funded Scheme (2018); Pearl River S&T Nova Program of Guangzhou (201806010183); Province Science and Technology of Guangdong Research Project (2017A020208055); Guangdong Key S&T Program (Grant no. 2019B020217002) from Department of Science and Technology of Guangdong Province; Walmart Foundation (SA1703162); National Broiler Industry Technology System Project (cARS-41-G16). The funders have no role in the study design, data collection and interpretation, or the decision to submit the work for publication.

## Declaration of complete interests

The authors have no conflicts of interest to declare.

## Ethic approval and consent of participate

This work only used bacterial strains and does not involve the use of human samples.

## Consent for publication

Not applicable.

## Footnotes

1 Evidence: On January 21, 2021, we searched PubMed with the terms “*Salmonella* Weltevreden”, “ST365”, “Human diarrhea”, and “Animal Diarrhea” for reports published, with no language restrictions. Our search identified no results of relevance to this study; we then searched PubMed with the terms “*Salmonella* Weltevreden ST365 in humans” for reports published, with no language restrictions. Two reports (PMID: 32983012 and PMID: 26496617) are listed but only one (PMID: 32983012; reference [8]) is associated with human diarrhea. In particular, none of the above studies reported the structure of the T4SS-carrying-IncFII(S) type plasmid and its association with the virulence of *S*. Weltevreden.

